# Quantitative Logics for Directed Evolution of an Asexual Population

**DOI:** 10.1101/2025.11.27.690992

**Authors:** Kangbien Park

## Abstract

Directed evolution of asexual populations is expected to offer a wide range of benefits to humanity. Achieving efficient directed evolution (DE) requires a quantitative formulation of the experimental methodologies—or logics—that can address potential challenges throughout the evolutionary process. In this article, I introduce ten logics designed for specific purposes in DE, accompanied by their mathematical formulations. To illustrate their application, I consider a hypothetical scenario of evolving an imaginary asexual population limb into a wing, employing a matrix-based discretization method to represent the trait of interest. The results indicate that applying these logics can accelerate the evolutionary process by roughly tenfold, while achieving an average accuracy of approximately 85% in reaching the desired trait across ten simulation iterations. Based on these findings, I discuss key considerations for implementing quantitative logic–based DE, propose strategies to improve alignment between the final outcome and the target trait, and examine factors for making the discretization method practical. Finally, I explore potential connections between quantitatively guided DE processes and emerging technologies, such as artificial intelligence and quantum computation.

## 2 Introduction

Directed evolution (DE) of asexual populations is an active area of research and is expected to bring diverse benefits to humanity Yuan et al. [2005], Laurent et al. [2023]. Despite substantial experimental progress in implementing DE through biological and chemical techniques Ren et al. [2025], Lee et al. [2020], quantitative modeling remains a crucial component of the DE process LaCroix et al. [2017], Lalejini et al. [2022], Corrao et al. [2024]. In particular, mathematical formulations enable rigorous analysis and the development of strategies to overcome key challenges associated with evolutionary experiments.

For example, when a population becomes trapped at an undesired local optimum on the fitness landscape, methods are needed to steer the population along pathways that descend the landscape. Rigorously achieving this requires precise mathematical modeling of both the trait and its associated potential. Additionally, if the evolutionary process takes too long to reach the target algebra, strategies for accelerating the trajectory become necessary. To this end, predictive tools that estimate the evolutionary timescale based on the mathematically defined experimental design, and that suggest shortcuts informed by such predictions, are essential.

Finally, mathematical formulations also provide insight into how changes in environmental conditions or differences in evolutionary speed influence the overall success probability of the evolutionary process, as these outcomes are ultimately governed by stochastic evolutionary dynamics.

Fortunately, with the growing body of knowledge in evolutionary dynamics and population genetics, along with the rapid increase in computational power driven by advances in computer science and AI, it has become increasingly promising to apply quantitative methodologies to directed evolution (DE) of asexual populations James et al. [2024], Gallup et al. [2024], Lee and Steel [2022]. Numerical approaches that design and simulate the evolutionary trajectory of an asexual population—and that express the corresponding experimental methodologies, or *logics*, in well-defined mathematical terms—can therefore serve as a foundation for systematically guiding and optimizing the DE process.

In this article, ten plausible logics for artificially directing evolution are mathematically formulated, and the corresponding reductions in evolutionary timescale are estimated. To accomplish this, an algebraic model based on an *n*_1_ *× n*_2_ *× n*_3_ matrix is used to represent the trait of interest, capturing the physiological structure of an imaginary asexual population limb in the *x*-, *y*-, and *z*-directions. This representation is constructed using a discretization method that converts a continuous geometric structure into a matrix-based form. The results show that these logics can accelerate the evolutionary process by an order of magnitude. In addition, the potential implementation of these logics in laboratory settings is discussed.

## 3 Theory and Methodology

For the overall analysis, an algebraic model Park [2025] based on the operator framework is employed Park and Bae [2024]. This model assumes an idealized situation in which mutations of interest occur only within the algebraic set, while the potential is defined independently of the current trait state. Note that I use the concept of the potential Φ = *F*_*max*_ − *F* where *F* is the fitness landscape Park [2025]. Under this setting, a two-dimensional vector with continuous components is used to provide an intuitive illustration of why the logics are important and how they can be defined.

Fig 1 illustrates the scheme for directing the evolution of a two-dimensional vector. The red arrows indicate the possible mutation directions within the plane, while the black arrows represent mutations in the *z*-direction (or any other higher-dimensional direction), which are forbidden under the assumption of a closed algebraic set. In this setting, the goal of DE is to guide the current population state (red ball) toward the target state (white ball), where the radius of each ball denotes the proportion of existing mutations.

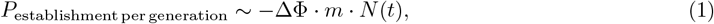

**Figure 1.**
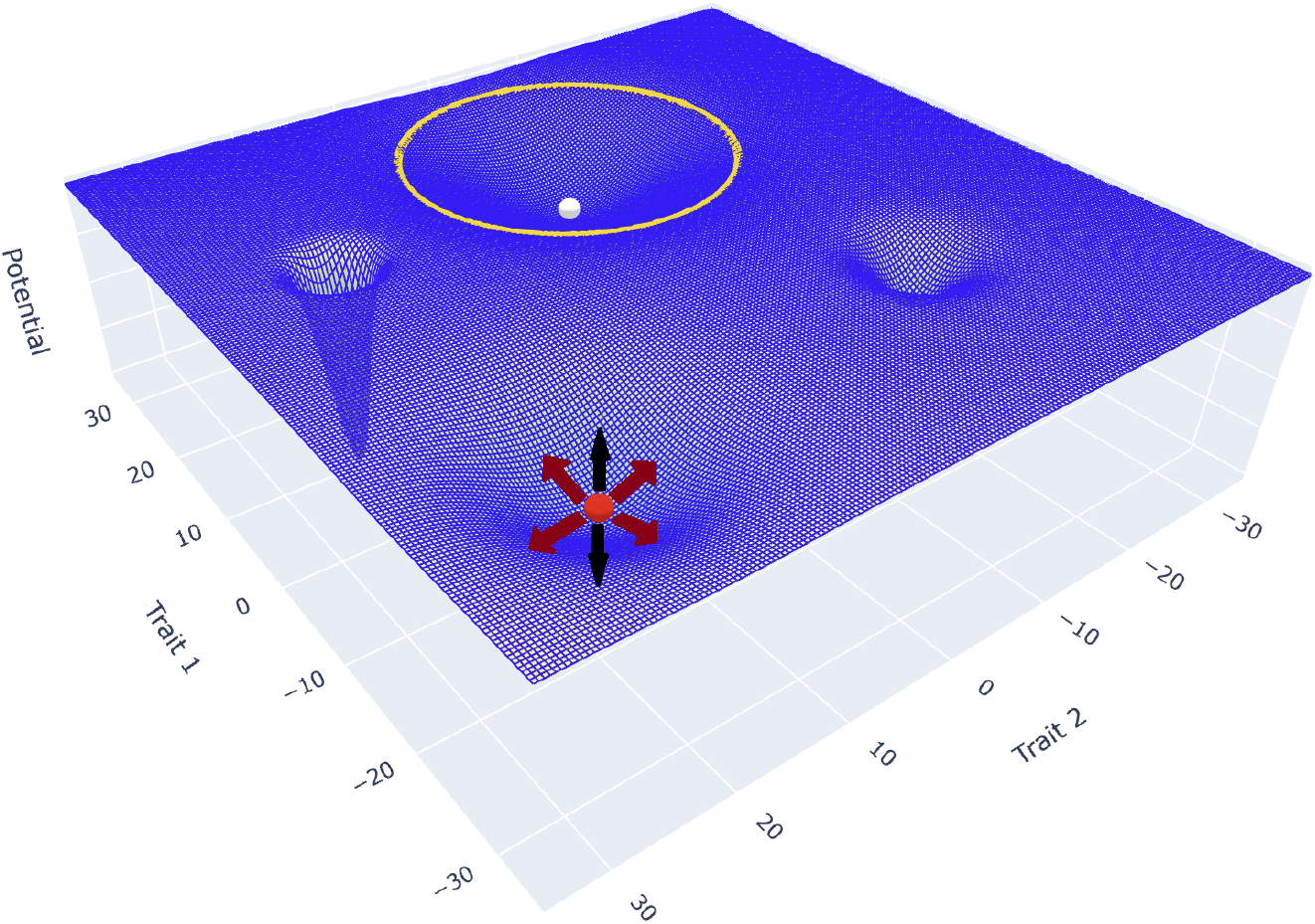
Illustration of the objective of directed evolution.

However, because the potential is convex near the red ball, artificial interventions—or *logics*—are required to push the ball upward along the potential Φ. Otherwise, an establishment event leading to a higher potential level will not occur naturally, since equation (1) implies that *P*_establishment per generation_ *<* 0 whenever ΔΦ *>* 0. Furthermore, even after the population reaches the flat region, additional guidance is required to reach the yellow ring. Once the population arrives at the ring, the evolutionary process naturally converges to the target position, as described by equation (2) [Park, 2025], where *A*_*i*_ represents the algebra of interest in the *i*-th generation, 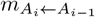 is the mutation rate from algebra *A*_*i*−1_ to *A*_*i*_, and 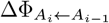 is the potential difference between *A*_*i*−1_ and *A*_*i*_.

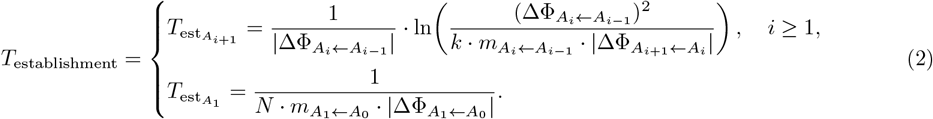

To achieve this, ten logics are introduced in this article: the logic of switching traits, the logic of towing from linkage, the logic of towing from phenotypic dependence, the logic of towing from pleiotropy, the logic of emergence, the logic of varying mutation rate, the logic of paste, the logic of positive potential, the logic of negative potential, and the logic of explicit modification. While other logics may also be devised to manipulate evolutionary trajectories, this article focuses on these ten.

Starting with the logic of switching traits, represented by Fig 2 (a) and (b), consider a particular trait represented by a two-dimensional vector that can adopt two different forms depending on the environmental pH. When the pH is neutral, the potential for the vector corresponds to form (a); when the pH is acidic, the potential shifts to form (b) because the trait adopts a different configuration. In this scenario, even if the state becomes trapped in an undesired local minimum under neutral pH, it can still evolve toward the target position when the environment turns acidic. The estimated time for the algebra to reach the objective position can be calculated using relation (2), with the potential differences derived from form (b).

**Figure 2.**
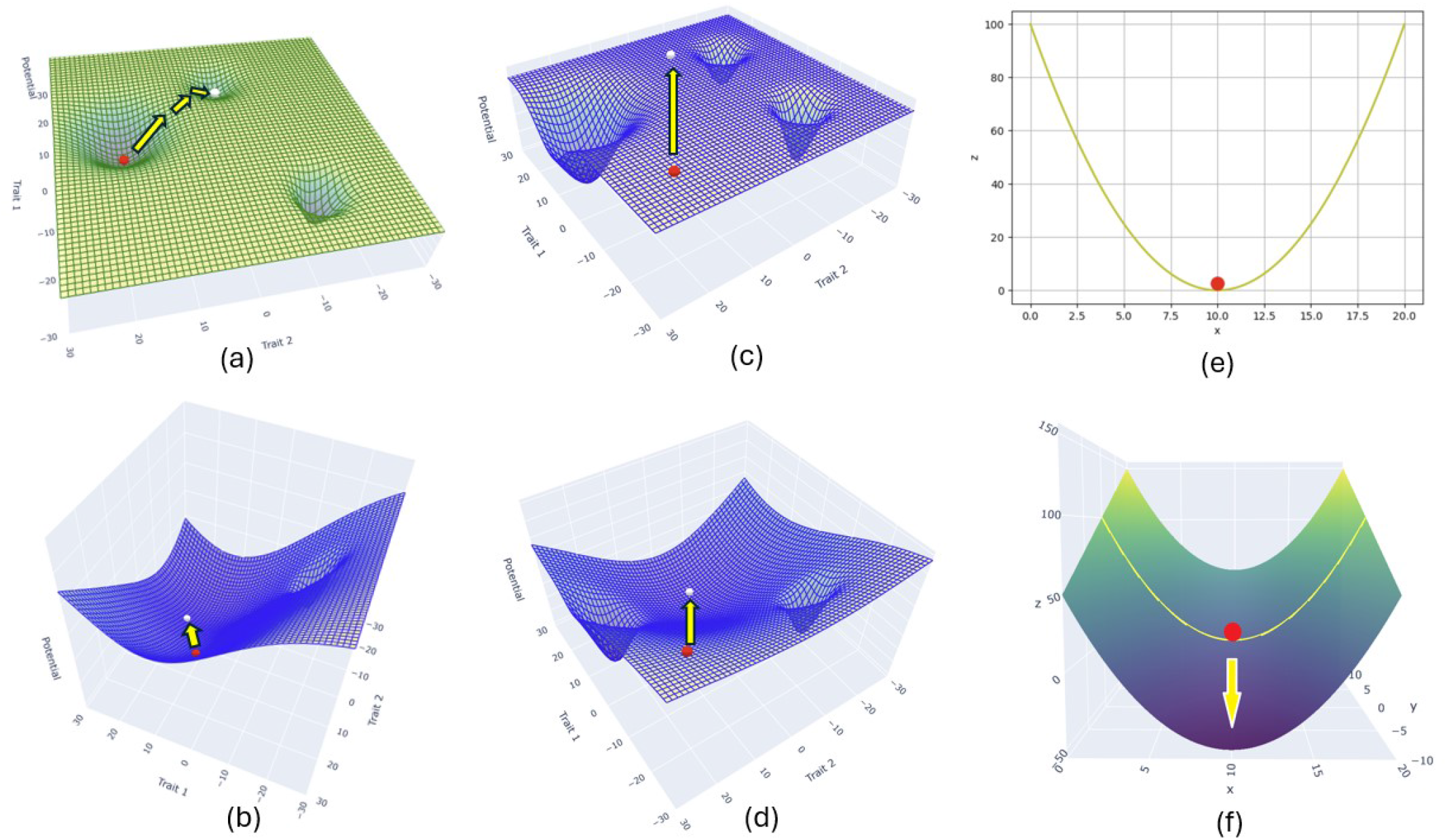
Pictorial representation for the logics with 2D vector case. (a) and (b) represent the logic of switching traits as well as the three variants of the towing logic. Also, (c) and (d) illustrate the logic of positive potential, and panels (e) and (f) demonstrate the logic of paste.

**Figure 2.**
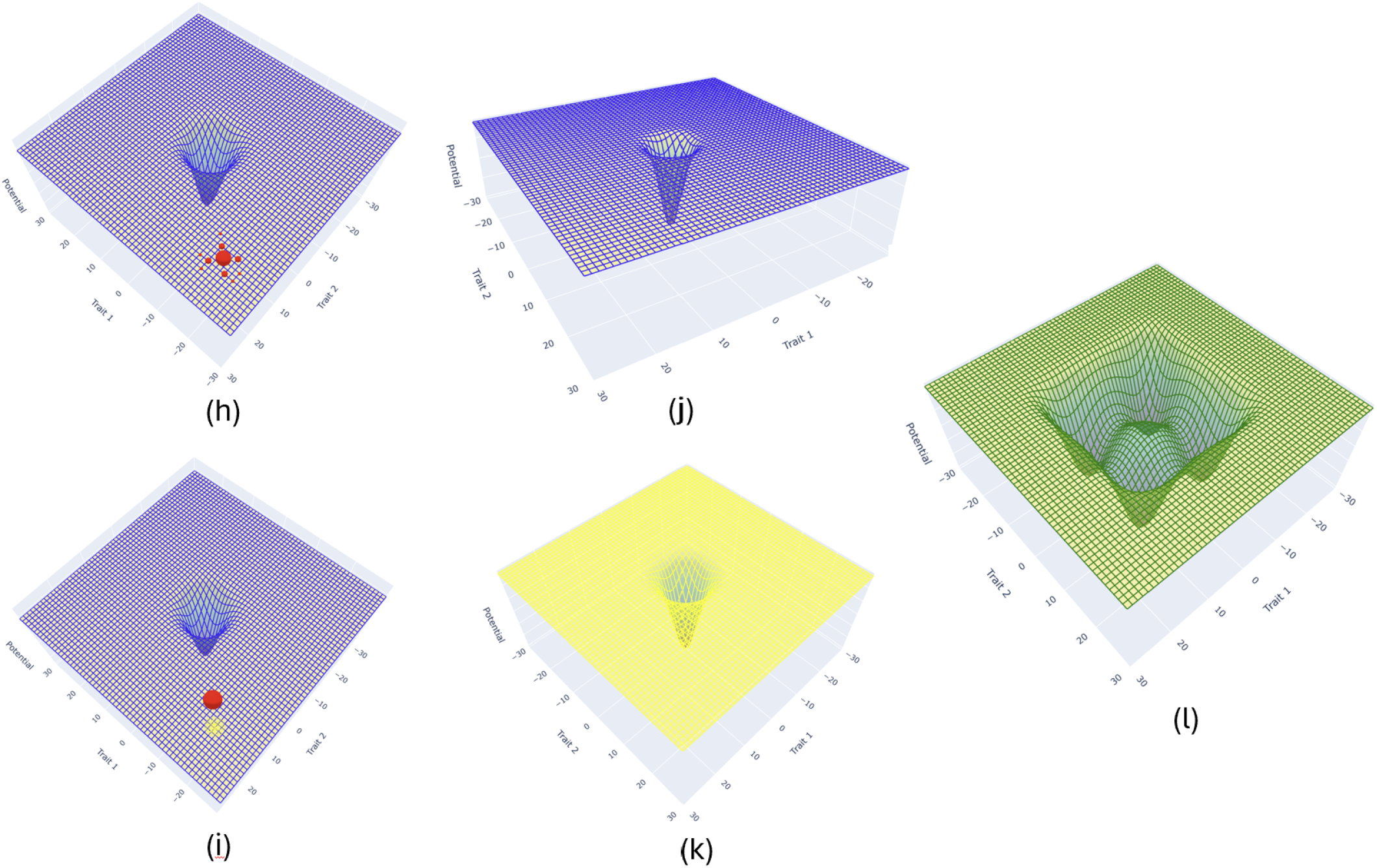
Continuous pictorial representation for the logics with 2D vector case. (h) and (i) represent the logic of negative potential. Also, (j), (k), and (l) illustrate the logic of emergence.

Second, the logic of positive potential, represented from Fig 2 (c) and (d) concerns altering the significance, or weight, of a particular environmental factor for survival, which induces an overall change in the potential. This change can be represented by form (c) as the initial potential and form (d) as the new potential after applying the logic of positive potential. The time required for the evolutionary reaction can then be estimated using relation (2), with the potential in this case taken from form (d).

Third, the logic of paste is essentially about expanding the explorable range of the algebra. For an intuitive example, consider an asexual population with only 100 base pairs of DNA, initially positioned at a minimum potential. By adding 100 more base pairs—analogous to “pasting” in sculpture—the population acquires additional genetic material, enabling further evolutionary exploration. In algebraic terms, the logic of paste typically involves increasing the dimension of the algebra. For instance, the initial algebra represented by a one-dimensional variable in Fig 2 (e) can be expanded into a two-dimensional space, as shown in Fig 2 (f). While no further optimization was possible in the original one-dimensional algebra, the logic of paste allows optimization in the higher-dimensional representation. Note that the logic of paste preserves all original algebra elements within the expanded set.

Fourth, the logic of towing from linkage relates to the evolution of linked genes, where multiple genes are physically located on the same plasmid or the main chromosome of the microorganism. For illustration of the logic, consider two genes encoding two 2-dimensional vectors that are linked within a plasmid. Even if the first vector should move downward along its potential, it may be constrained if the selective advantage of the second vector is much larger, preventing the first vector from moving as expected. In this scenario, even if the first vector becomes trapped in a local minimum, as illustrated in Fig 2 (a), mutations corresponding to a higher potential can still establish due to the dominant evolutionary influence of the linked second vector. It is important to note that this approach can also stabilize the position of the first vector, effectively *desensitizing* it to potential differences because of the dominant dynamics of the second vector.

Fifth, the logic of towing from phenotypic dependence concerns two interdependent algebras, which do not necessarily share genes. In this scenario, the evolution of the first algebra can alter the potential of the second algebra. While generation-independent potentials are generally preferable for ease of calculation, this logic requires accounting for the generation-dependent nature of the potential. To simplify the analysis, one could experimentally restrain the evolution of the second algebra until the first algebra completes its influence on the second algebra’s potential, and then allow evolution to proceed. Under this approach, the total generation for the evolutionary reaction would include the additional duration required for the potential change.

Sixth, the logic of towing from pleiotropy exploits the fact that a single gene can influence multiple algebras. This logic is structurally similar to the logic of linked genes, where the dominant algebra determines the evolutionary trajectory of a less strongly selected algebra. However, unlike the linked case, the effect in pleiotropy is more direct: the weaker-selected algebra must move whenever the dominant algebra moves. The evolutionary time in this scenario can be estimated by considering the dominant algebra’s evolution via relation (2) and its effect on the DNA sequence, which subsequently drives the evolution of the target algebra. For precise modeling, the relationship between the DNA sequence and the algebraic representation of the corresponding phenotypes must be clearly defined.

Note that the three variants of the logic of towing are illustrated across Fig 2 (a) and (b): Fig 2 (a) depicts the primary algebra of interest, while (b) shows the secondary algebra that facilitates towing.

Seventh, the logic of negative potential is inspired by the behavior of antibiotic-resistant bacteria. When a mutation allows survival in a harsh environment, it is almost certainly fixed, regardless of whether it was initially established. This harsh environment is represented as a negative potential, illustrated in (h) and (i) of Fig 2. In (h), the radius of the ball represents the proportion of the mutation, showing that only a few vectors occupy the position (-16, 16). After applying the negative potential, which permits survival only at (-16, 16), the original population around (-20, 20), shown by the dotted yellow sphere, is driven to (-16, 16), illustrated by the solid red sphere. The benefits of the negative potential logic are twofold. First, it accelerates evolution by bypassing the time required for beneficial mutations to be established and fixed. Second, in directed evolution, key intermediate states are often needed but may be difficult to establish. In such cases, negative potential can eliminate competing variants, leaving only the algebra of interest and thus contributing to the overall success of the evolutionary reaction. The total evolutionary time can then be estimated as *t* = log_2_(*N/N*_initial_), representing the time required for the surviving mutation population to double from *N*_initial_.

Eighth, the logic of emergence concerns the combination of potentials that produces emergent effects. It is often assumed that when two potentials, Φ_1_ and Φ_2_, have the forms shown in Fig 2 (j) and (k), their sum, Φ_sum_, can be represented simply as a linear addition, Φ_sum_ = Φ_1_ +Φ_2_. However, some potentials may exhibit emergent properties, such that Φ_sum_ = Φ_1_+Φ_2_+*f*(Φ_1_, Φ_2_), where *f*(Φ_1_, Φ_2_) is an emergent function, resulting in the form illustrated in (l). To apply the logic of emergence in directed evolution, extensive experimental data are required to accurately combine the potentials and design the emergent potential. The expected evolutionary time can then be estimated using relation (2), with the emergent potential taken into account.

Ninth, the logic of varying the mutation rate concerns adjusting the mutation rate of the population. Since the mutation rate strongly influences the evolutionary trajectory, strategically increasing or decreasing it at specific stages can help guide the population along the desired path more efficiently.

Tenth, the logic of explicit modification involves directly moving the red ball to or near the objective position. In practice, this can be realized in the laboratory through genetic engineering methods, effectively placing the population closer to the desired trait.

Using the introduced logics, I evolve the traits of interest represented by an *n*_1_ *× n*_2_ *× n*_3_ matrix *M*, demonstrating an application of algebraic modeling to directed evolution (DE) [Park, 2025]. This matrix contains only 0s and 1s, where 1s correspond to points defining the physical shape in three-dimensional space. Rather than using the DNA sequence as the basis for mutation, mutations are applied directly to the algebra *M* representing the phenotypic trait, such that an element of *M* changes from 0 to 1 after mutation. Mathematically, the algebra is defined as

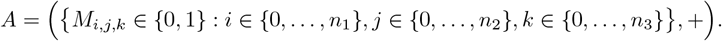

Although other algebraic representations could be used to model the traits of interest, this *n*_1_ *× n*_2_ *× n*_3_ matrix was chosen to intuitively capture the continuous physiological geometry of the traits in three-dimensional space via discretization, as illustrated in Fig 3.

**Figure 3.**
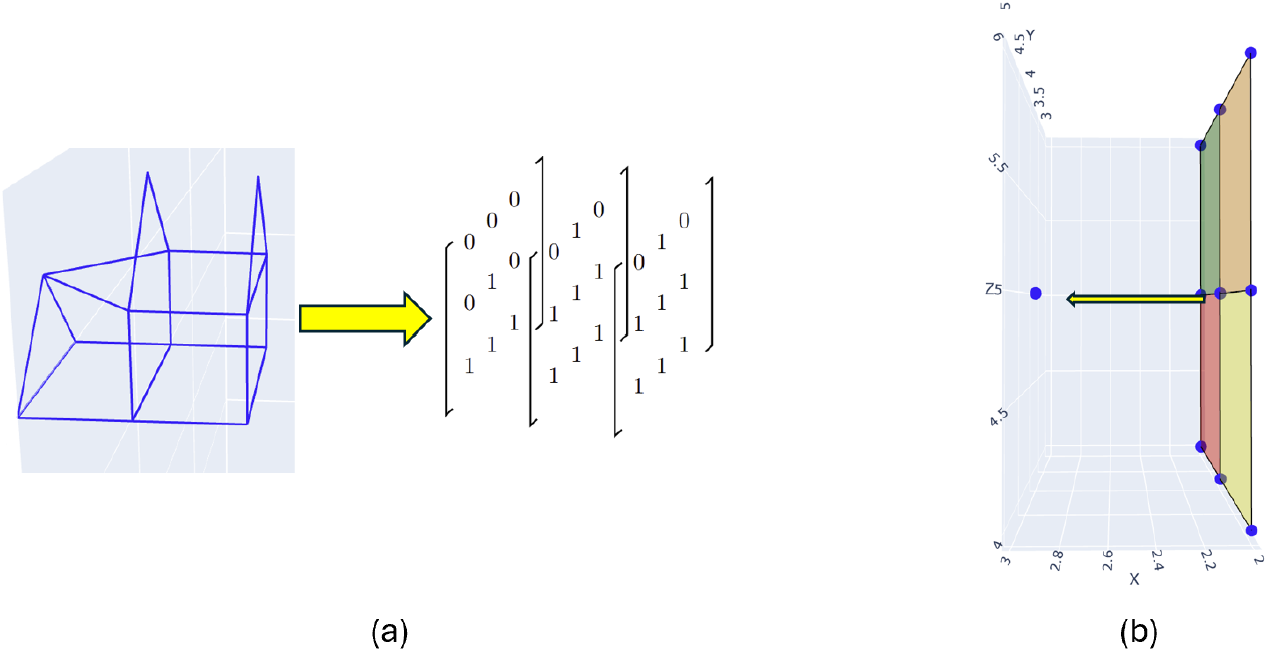
(a) illustrates the discretization example, while (b) represents the square-based mutation rule.

**Figure 4.**
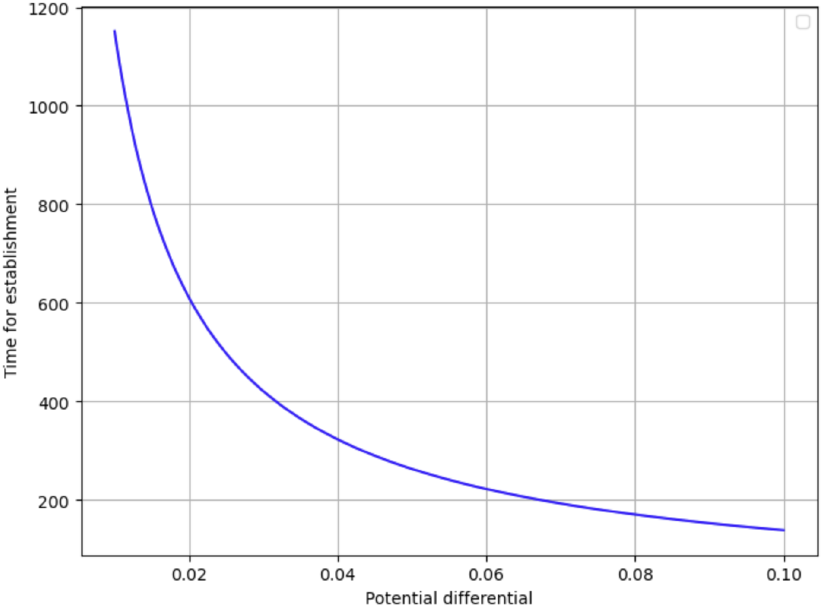
Graph representing the relation between ΔΦ and time taken for establishment.

With the defined algebra, consider a scenario in which the limb of an imaginary asexual population—sharing evolutionary characteristics similar to bacteria—gradually evolves into a wing. For this process, research on the evolutionary development of wings in birds, insects, and bats was used to define the appropriate potentials. Specifically, the development of the patagium enabling thermoregulation [Kingsolver and Koehl, 1985, Shockney and Rummel, 2025], the semi-wing structure allowing proto-birds to generate thrust for climbing trees [Bundle and Dial, 2003], and the wing structure enabling lift for flight [Tobalske, 2007] were all considered.

For the matrix, the mutation law is defined as follows: a mutation occurs when an original element 0 turns into 1. This mutation occurs only if there is an existing background structure in the direction of the mutation. To illustrate with an example, when the matrix evolves in the *x* direction at position *M*_3,4,5_, one of the following sets of elements must already be 1:

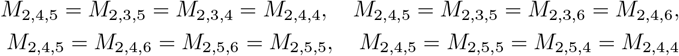

as illustrated in Fig 3 (b). While other mutation rules could be applied, this choice intuitively represents a scenario in which a new mutation occurs only when there is sufficient prior “paste” in the matrix.

Now, with a well-defined matrix and its mutation law, the evolutionary path depends on the geometry of the potentials. To list the potentials that are used, first the thermal regulation potential is approximately modeled as

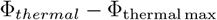

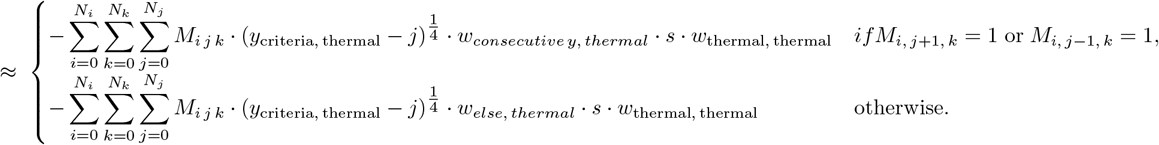

The second potential, the thrust potential, is defined as

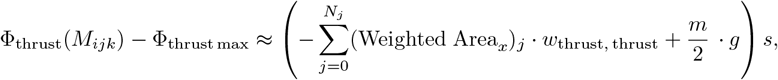

where the weighted area is the *x*-plane projected area computed according to the following rule:

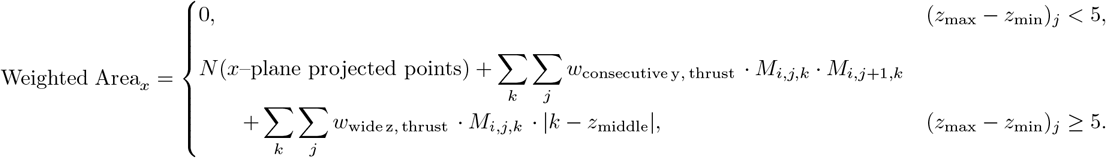

In detail, the weighted area is computed as follows. First, the total number of *x*-plane projected points is counted. Then, for each pair of consecutive *y*-values at the same *z*-value, a weight *w*_*y*_ is added, and for each integer *z*-value, a weight *w*_*z*_ · (*z* − *z*_middle_) is also summed, where *z*_middle_ denotes the midpoint index of the matrix along the *z*-axis. These considerations are introduced to approximately capture the action–reaction principle for real gases: when the wing is insufficiently broad, the gas tends to slip sideways rather than exerting thrust.

Moreover, the mass *m* is defined to be proportional to the total number of matrix elements, given by

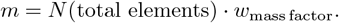

Next, the lift potential is defined as

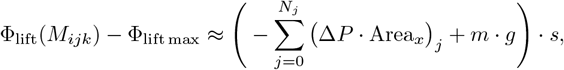

where the pressure difference for each *j* is given by

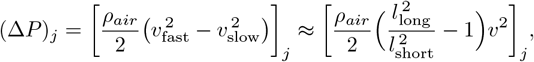

following Bernoulli’s principle, based on the wing’s y-axis cross-sectional structure shown in Fig 5 (h). Here, *l*_long_ and *l*_short_ denote the longer and shorter perimeters of the wing, respectively, and *v* is the velocity of the moving creature, approximated as *v* ≈ *w*_velocity, lift_. Moreover, *ρ*_*air*_ = 1.225 (omitting an explicit physical unit *kg/m*^3^) is set to represent the realistic value of the air density.

**Figure 5.**
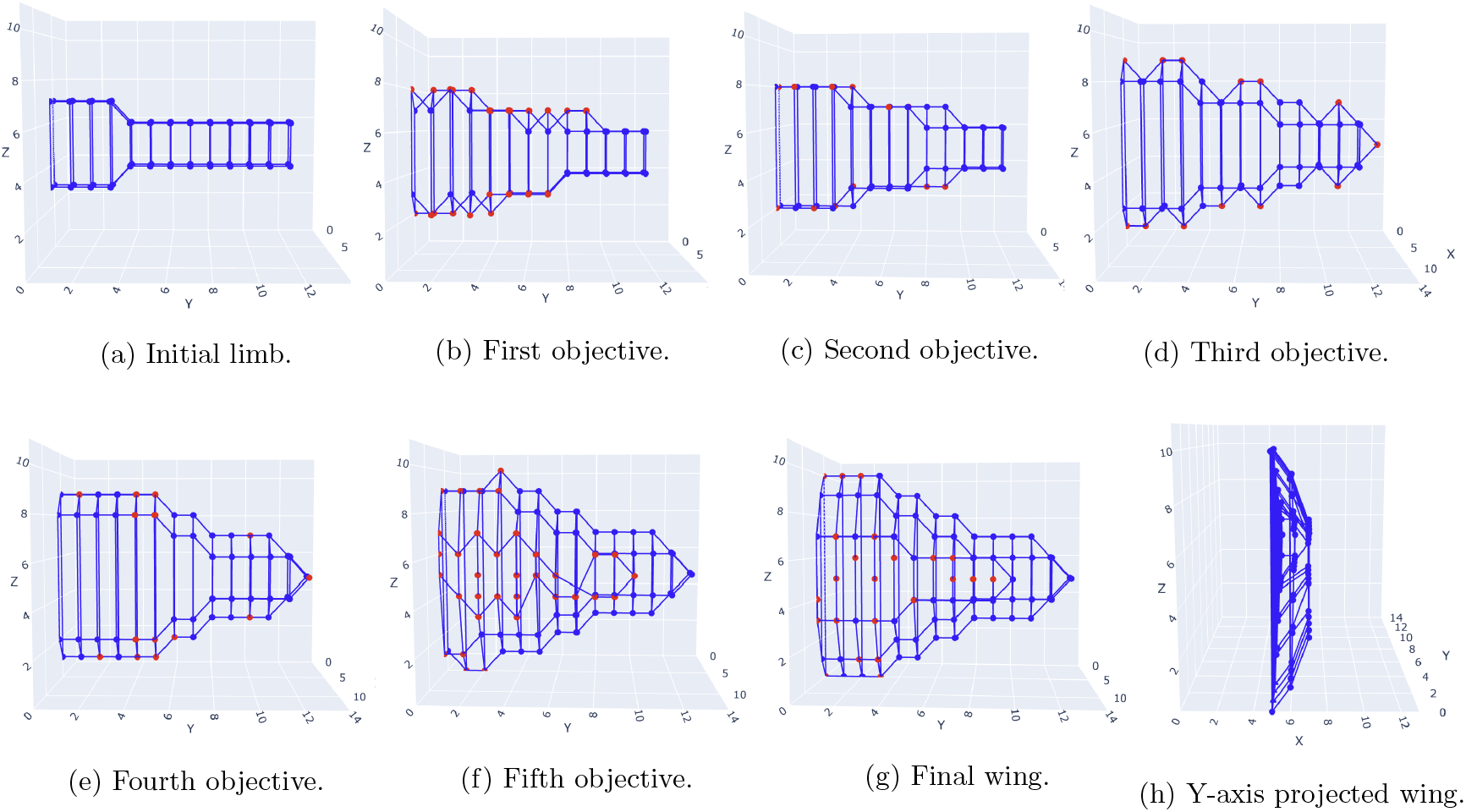
The graphical representation of milestones in the directed evolution of the discretized matrix. The red dots indicate elements that evolved from the preceding objective.

The quantity Area_*x*_ is the *x*-plane projected area for each *j*, obtained by projecting all matrix elements with *y* = *j* onto the *x*-plane and counting the number of projected points. The following constraints are applied. First, if the wing is too thick at the edge-specifically, if Δ*x* ≥ 3 for a fixed *y* and for either *z*_min_ or *z*_max_-then that *y*-plane component does not contribute to the lift force, based on the assumption that excessively thick edges prevent the airflow from splitting cleanly enough to generate Bernoulli lift. Second, if the angle between adjacent points within a fixed *y*-plane exceeds 45^°^, the corresponding component is also excluded from contributing to lift, under the intuitive assumption that airflow may not smoothly follow such abrupt geometric changes.

Lastly, the towing potential is modeled in a straightforward manner, taken to be proportional to the total number of matrix elements that exceed a specified threshold in the *x, y*, or *z* direction:

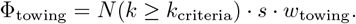

The emergent potential is similarly modeled in a simplified but nonlinear form:

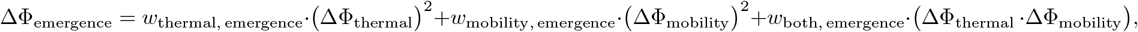

where the weight vector is defined as

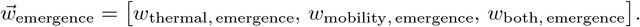

Within this potential framework, several observations can be made. When only the thermal potential Φ_thermal_ is active, the optimal matrix configuration is one in which all elements with indices satisfying *j* ≤ *y*_criteria, thermal_ are equal to 1. When only the thrust potential Φ_thrust_ is applied, multiple optimal configurations exist, each maximizing the *x*-plane projected area. Finally, when the lift potential Φ_lift_ is the only active potential, the matrix evolves toward a configuration that maximizes the ratio between *l*_long_ and *l*_short_.

Importantly, these optimal matrices differ substantially from one another, underscoring the necessity of adjusting the activated potentials at appropriate moments to guide the system toward the target matrix shown in Fig 3 (b).

The total potential combining these effects is defined as

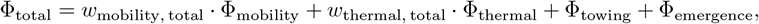

where

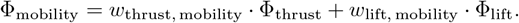

The table 1 shows the initial weight values that determine the total potential. These weights are experimentally adjustable parameters, tuned according to the quantitative logics introduced above. They are set so that each mutation approximately lowers the potential by ΔΦ ≈ −0.01 ∼ −0.1, with an average value of ⟨ΔΦ ⟩ ≈ − 0.02, consistent with empirical observations of natural evolution [Imhof and Schlötterer, 2001]. Under these conditions, each newly established mutation requires approximately *T* ≈ 841 generations according to equation (2), assuming a mutation rate of *m* ∼ 10^−9^ [Imhof and Schlötterer, 2001] and *k* ≈ 1 [Park, 2025]. Now, the initial limb shown in Fig 5 (a) contains 94 elements equal to 1, whereas the final wing in Fig 5 (b) contains 214 such elements. Thus, a rough estimate indicates that the transition requires approximately 214 − 94 = 120 mutational establishments, corresponding to about 120 *×* 841 ∼ 10^5^ generations.

**Table 1:** Weights are set such that mutations can decrease the potential on the order of ΔΦ ∼ −0.02.

| Weight | Value |
| --- | --- |
| $w_{\text{mass factor}}$ | 1/50 |
| $g$ | 10 |
| $w_{\text{thrust, mobility}}$ | 1/5 |
| $w_{\text{lift, mobility}}$ | 1/40 |
| $w_{\text{thermal, total}}$ | 1/3 |
| $w_{\text{mobility, total}}$ | 2/3 |
| $y_{\text{criteria, thermal}}$ | 10 |
| $w_{\text{thermal, thermal}}$ | 1/2 |
| $w_{\text{consecutive } y, \text{ thermal}}$ | 10 |
| $w_{\text{else, thermal}}$ | 5 |
| $w_{\text{thrust, thrust}}$ | 12.5 |
| $w_{\text{consecutive } y, \text{ thrust}}$ | 1/5 |
| $w_{\text{wide } z, \text{ thrust}}$ | 1/10 |
| $w_{\text{velocity, lift}}$ | 1 |
| $w_{\text{towing, } x}$ | 0 |
| $w_{\text{towing, } y}$ | 0 |
| $w_{\text{towing, } z}$ | 0 |
| $\vec{w}_{\text{emergence}}$ | $\vec{0}$ |

Even assuming a generation time of roughly 30 minutes for an asexual population, the entire process would then require about 70 months, making it economically impractical. Furthermore, the evolutionary trajectory is not guaranteed to reach the target matrix, since many alternative mutational paths exist and evolutionary outcomes are irreversible.

To overcome such inefficiency in the number of required generations, the logics introduced above can be employed to design the evolutionary pathway. In doing so, it is important to recognize several fundamental principles. First, the matrix is more likely to evolve in the direction that maximally decreases the potential, i.e., along −∇ Φ, as implied by equation (1). Second, to minimize the reaction time, the evolutionary path should ideally follow a trajectory of roughly uniform potential decrease [Park, 2025], unless the logics provide novel shortcuts. Based on these principles, the evolutionary pathway guided by the logics is designed to achieve potential decreases of ΔΦ ≈ − 0.1 ∼ − 0.4. Such a setting could be experimentally feasible given the relatively strong selection that bacteria can experience [Sidorova et al., 2020]. Moreover, a beneficial mutation rate of *m* = 10^−7^ is chosen, as it falls between previously reported empirical values—ranging from *m* ∼ 10^−9^ [Imhof and Schlötterer, 2001] to *m* ∼ 10^−5^ [Perfeito et al., 2007]—and thus represents a realistic and experimentally attainable value. With these parameters, the estimated evolutionary time decreases to approximately 3,000–16,500 generations, corresponding to roughly 3–11.5 months, which is about one-tenth or less of the time required under natural evolutionary conditions, while the accuracy of reaching the objective is also expected to improve.

Note that the population evolution in this setting appears to deviate from the power-law behavior of selection coefficient accumulation, even though the environment is time-independently fixed [Wiser et al., 2013], and instead exhibits an approximately linear accumulation of selection coefficients. Despite this seemingly apparent discrepancy, the evolution still fundamentally follows the power law; the observed linear behavior corresponds only to the initial range of the power-law curve. Over a much larger number of generations, the matrix evolution will eventually converge to the expected power-law trend.

The evolutionary path is illustrated in Fig 5. It begins by allowing thrust force to be the dominant environmental pressure ((*a*) → (*b*), 3,000 generations), followed by thermal regulation becoming the primary selective factor ((*b*) → (*c*), 2,000 generations). These pressures are then repeated for one more cycle ((*c*) → (*d*), 1,500 generations; (*d*) → (*e*), 1,500 generations), before the environment increasingly requires lift force ((*e*) → (*f*) → (*g*), 5,000 generations). To realize such qualitative design, the logic of towing and logic of positive potential were actively used while the logic of negative potential was often used. To achieve this, quantitative parameters were adjusted to approximately maximize the potential drop along the desired mutation direction, thereby increasing the probability of establishment in that direction. The specific weights used are provided in the Appendix.

For the simulation, all logics related to towing were incorporated into the logic of towing defined above, while the logic of switching traits was included in the logic of positive potential. The logics of explicit modification, varying mutation rate, paste, and emergence were not applied in this instance, as they were unnecessary for evolving the wing. However, these options remain available and can be simulated using the posted code. Note that the simulation was developed based on the operator model [Park and Bae, 2024], and ChatGPT was utilized to assist in developing sections of the plotting code and to support the design and debugging of specific parts of the computational elements.

## 4 Results and Discussion

The simulation results for a 13,000-generation evolution followed the predesigned trajectory, with the target wing structure shown in Fig 5 (g) (the resulting wing morphologies are provided in the appendix). The matrix corresponding to this figure was selected as the reference final matrix among several plausible divergent structures because it preserved the key morphological features intended to define a wing. Moreover, the morphology in (g) aligns with a rough quantitative expectation: if the weights are set such that commonly observed new mutations produce a potential change of approximately ΔΦ ≈ − 0.15, then relation (2) predicts that the final structure would incorporate about 120 new points over roughly 13,000 generations. Therefore, the matrix represented by Fig 5 (g) serves as an informed reference that both reflects the desired wing shape and aligns with predictions from a rough calculation. Moreover, to refine this reference, preliminary iterations were carried out to predict the final wing shape under weight sets that typically produce mutations with ΔΦ ∼ − 0.1 to − 0.4 and steer the evolution toward a morphology resembling Fig 5 (g). The outcomes of these preliminary simulations were then used to estimate the expected final wing structure, ultimately supporting the choice of Fig 5 (g) as the representative form. Although this process of performing preliminary iterations and anticipating the final form may seem somewhat ad hoc, it illustrates a realistic approach that computational DE can follow—namely, an expectation-maximization (EM) algorithm [Dempster et al., 1977]. Specifically, if the experimenter identifies a desired trait with certain degrees of freedom, adjusting the weights can help constrain that freedom and identify the optimal solutions among the diverse candidates sharing the trait. This is particularly important because, in realistic scenarios, objective traits often allow a wide range of functional or morphological variations that cannot be fully specified by the experimenter’s criteria.

The results in Table 2 demonstrate that an informed approximation of the objective was indeed feasible, and that the matrix can evolve into traits resembling the target characteristics. In particular, the resulting wings recovered, on average, up to 85% of the objective points, and the total number of 1s differed only slightly from the expected values. Table 2 also suggests that, for the system to approach the intended morphology, the evolutionary pressures associated with lift and thrust should gradually decrease, whereas the pressure associated with thermal regulation should increase.

**Table 2:** Quantities were measured from 10 independently gevolved wings, each obtained through a separate iteration of the procedure. The term “included points” refers to the percentage of points associated with the objective trait that appear in the resulting matrix, whereas the “total number of 1s” denotes the percentage difference between the total count of 1s in the resulting matrix and that in the objective matrix. Percentage differences for the potential values of lift force, thermal regulation, and thrust force were calculated using the weights listed in Table A3, which correspond to the environmental conditions at the final stage of the evolutionary process. Here, the percentage difference was computed using the formula 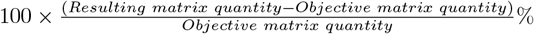.

| Comparing Quantity | Min (%) | Max (%) | Average (%) |
| --- | --- | --- | --- |
| Included Points | 80 | 89 | 85 |
| Total Number of 1s | -6 | 9 | 2 |
| Lift Force | 11 | 52 | 29 |
| Thrust Force | -2 | 20 | 9 |
| Thermal Regulation | -24 | 8 | -5 |

Despite this, the outcome does not perfectly match the desired objective for two principal reasons: the limited computational capability to precisely track the evolutionary trajectory, and the restricted number of variables available for optimization. If the simulation were designed to evaluate potential changes for all plausible mutations at each generation of the directed evolution process—while simultaneously adjusting the associated potentials using optimized weights—the accuracy with which the target trait is achieved would improve substantially. Such an enhanced system, capable of handling large-scale calculations, would enable the design of far more sophisticated evolutionary pathways. Nonetheless, the primary goal of this work was to demonstrate that directed evolution guided by quantitative principles is feasible. Accordingly, the implemented code was developed only to the extent needed for this proof of concept.

Moreover, controlling only the three potentials listed above imposes limitations on achieving the exact target matrix. This is because multiple divergent evolutionary pathways exist, and in many cases, undesired traits can yield a similar or even greater potential decrease than the objective trait under the preset weights. For instance, the symmetry evident in Fig 5 (g) cannot be easily ensured, as the listed potentials do not regulate the additional degrees of freedom required to produce a symmetric wing geometry. Similarly, evolution along the *x*- and *z*-axes was difficult to control with the given parameters. This is because preparing combinations of parameters that maintain ΔΦ between − 0.1 and − 0.4 in the desired direction required selecting settings that were among the more persuasive options available (i.e., the weights listed in the appendix), yet these settings still allowed deviations along the *x*- and *z*-directions. In summary, sophisticated computational tracking and additional potentials that provide more degrees of freedom are essential for higher-precision DE, and would further enhance the accuracy of the resulting outcomes.

Also, even when the |ΔΦ| is fairly large (e.g., around 0.3 ∼ 0.4) for mutations, the evolutionary reaction still requires a minimum amount of time for any mutation to become established. Consequently, if the objective traits must go through many such establishment steps from the original traits, a significant minimum time is inevitably required. To reduce this time, one approach is to begin with traits already available in nature that are closest to the desired objective traits, or to utilize the logic of explicit modification. In such a genetic engineering approach, providing only a rough framework for the desired trait is sufficient, as it still allows numerous possibilities to evolve into the fully developed desired trait. On the other hand, when the experimenter cannot fully predict how the engineered genes will behave due to numerous emergent effects within the biological system [Zhang et al., 2016, Şimşek et al., 2023], DE methodologies can be employed to refine and stabilize the inserted genes so that they function as intended. Thus, genetic engineering and directed evolution can complement each other, working synergistically to achieve the desired traits.

As demonstrated, evolutionary modeling based on trait-geometry discretization can provide significant benefits for DE calculations. To apply a discretized trait matrix in practical DE experiments, it is essential to establish a well-defined mutation law that governs how the matrix evolves from DNA mutations. Ideally, this could be implemented within a continuous *in vivo* DNA mutation system [Tong et al., 2025, DeBenedictis et al., 2022, Wong et al., 2018, Steel et al., 2020], in which the evolution of matrix elements follows predictable rules.

One possible strategy is to introduce a *parallelly evolvable genetic system*—a combination of DNA or RNA sequences with regulatory pathways—that satisfies three major characteristics: orthogonality and modularity [Wang et al., 2011, Szenk et al., 2020] of gene elements, inducibility [Tominaga et al., 2024, Cui et al., 2016, Wiechert et al., 2020] of the control system, and malleability (or evolvability) [Pines et al., 2017, Wagner, 2023] of genes.

Specifically, orthogonality ensures that multiple, mutually independent DNA sequences collectively contribute to the formation of the discretized matrix. A key advantage of this design is that, by avoiding long serial DNA constructs, mutations are less dependent on preceding sequences, allowing greater flexibility without compromising the integrity of already-evolved segments. This modularity also helps prevent unwanted epistasis among elements. Regarding inducibility, if a DNA fragment in the parallel system remains dormant until specific combinations of other elements are active, this behavior may serve as a robust foundation for implementing a matrix-based discretized evolution framework. Finally, gene malleability ensures that DNA or RNA sequences can be readily modified under evolutionary pressure. In summary, employing a parallelly evolvable genetic system is analogous to introducing independent layers of malleable clay: initially unstructured, these layers are progressively shaped through evolutionary processes, while a discretized matrix framework constrains and guides the system’s evolutionary dynamics. With a genome engineered to satisfy the characteristics outlined above, it becomes possible to envision a framework for *standardized directed evolution* (SDE) system, in which the evolutionary behavior of phenotypic traits is quantitatively tractable. To rigorously develop such an SDE system, insights from genotype–phenotype mapping [Otwinowski and Nemenman, 2013, Ascensao and Desai, 2025] will be essential for the rational design and optimization of this approach. By eliminating the need for instance-specific analyses of individual asexual populations, an SDE framework is expected to provide a systematic and generalizable foundation for future directed evolution methodologies.

Moreover, the example above treated the traits defining the algebra as the spatial physiological structure along the x, y, and z directions, which allowed efficient calculation of physical potentials since both classical mechanics and the trait space could be naturally formulated in three-dimensional physical space. However, when considering more general traits in arbitrary mathematical spaces, it becomes necessary to develop both theoretical and empirical methodologies to define a well-posed map that links matrix elements to the corresponding potential. In summary, theoretical research aimed at devising abstract mathematical geometries that efficiently model traits of interest—while ensuring that mutations of the geometry remain predictable based on preset genome—would be of critical importance.

Finally, note that since the design of the evolutionary path is highly sensitive to even small changes in the potentials, a rigorous formulation that accurately reflects realistic environmental effects is imperative. In practice, because the potentials continuously change as the trait evolves, they will likely need to be remeasured at regular intervals during the evolution. Additionally, to achieve the required level of precision in potential measurement, massive datasets from both theoretical calculations and experiments are necessary, and the mechanisms by which environmental factors influence the potential must be well understood.

With such data in hand, the remaining task becomes the tuning of weights to design an evolutionary path that maximizes the success rate while minimizing the time required to reach the objective algebra using the logics described above. For the purpose, artificial intelligence (AI) methodologies, such as gradient descent [Cauchy et al., 1847] and backpropagation [Rumelhart et al., 1986], can be effectively employed to adjust parameters and minimize deviations from the intended path. Similarly, if sufficient data exists on how environmental factor determines the weights, the DE process could potentially be automated via AI-driven computation. Here, since the Expectation–Maximization algorithm has a close conceptual connection with self-supervised learning [Zhao et al., 2018, Si et al., 2010], situations involving a large number of possible evolutionary paths toward an initially vague final trait can benefit from self-supervised approaches, which help converge on the desired trait while simultaneously adjusting the weights. Furthermore, for optimizations involving very high-dimensional potentials, techniques such as quantum annealing [Finnila et al., 1994, Kadowaki and Nishimori, 1998] or QAOA [Peruzzo et al., 2014] from future fault-tolerant quantum computing frameworks [Aharonov and Ben-Or, 1997] could be useful. Here, the mathematical structure of population states distributed across diverse potential positions may allow leveraging the superposition property of qubits to facilitate quantum optimization.

## 5 Conclusion

From the quantitative modeling of the logics corresponding to diverse experimental methodologies for addressing issues in directed evolution (DE), it is possible to calculate how the evolutionary path can be more effectively designed. Using a discretized matrix representing limb geometry, along with well-defined mutation rules and physical potentials, it was observed that these logics indeed allow the DE process to proceed more efficiently. Specifically, the speed of evolution increased by nearly tenfold, while the accuracy of the final trait relative to the objective reached approximately 85% on average. Encouraged by these results, further advancement in the formulation of quantitative logics for DE, or a more rigorous conceptualization of trait discretization, may provide deeper insights into how DE should be conducted. Moreover, such a mathematical representation of DE is expected to benefit from emerging technologies, including artificial intelligence and quantum computing, in various applications.

## 6 Data availability

The code for the simulation could be found from https://github.com/TheDEnotes/Quantitative-Logics-for-DE.

## 7 Acknowledgments

This work was largely conducted at the Department of Physics, Yonsei University, and at DAMTP, University of Cambridge. I am grateful for the calm and encouraging environments offered by both institutions. Furthermore, ChatGPT was utilized to help improve the clarity and readability of the English text.

## 10 Appendix

**Table A1:**
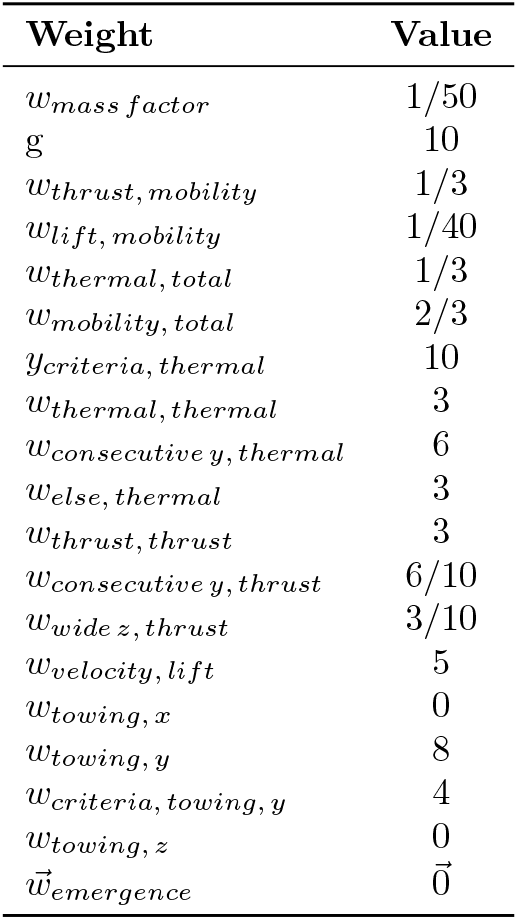
Weights for the thrust force step.

| Weight | Value |
| --- | --- |
| $w_{mass\ factor}$ | 1/50 |
| $g$ | 10 |
| $w_{thrust, mobility}$ | 1/3 |
| $w_{lift, mobility}$ | 1/40 |
| $w_{thermal, total}$ | 1/3 |
| $w_{mobility, total}$ | 2/3 |
| $y_{criteria, thermal}$ | 10 |
| $w_{thermal, thermal}$ | 3 |
| $w_{consecutive\ y, thermal}$ | 6 |
| $w_{else, thermal}$ | 3 |
| $w_{thrust, thrust}$ | 3 |
| $w_{consecutive\ y, thrust}$ | 6/10 |
| $w_{wide\ z, thrust}$ | 3/10 |
| $w_{velocity, lift}$ | 5 |
| $w_{towing, x}$ | 0 |
| $w_{towing, y}$ | 8 |
| $w_{criteria, towing, y}$ | 4 |
| $w_{towing, z}$ | 0 |
| $\vec{w}_{emergence}$ | $\vec{0}$ |

**Table A2:** Weights for the thermal regulation step.

| Weight | Value |
| --- | --- |
| $w_{mass\ factor}$ | 1/50 |
| $g$ | 10 |
| $w_{thrust, mobility}$ | 1/12 |
| $w_{lift, mobility}$ | 1/40 |
| $w_{thermal, total}$ | 1/3 |
| $w_{mobility, total}$ | 2/3 |
| $y_{criteria, thermal}$ | 10 |
| $w_{thermal, thermal}$ | 4 |
| $w_{consecutive\ y, thermal}$ | 10 |
| $w_{else, thermal}$ | 5 |
| $w_{thrust, thrust}$ | 3 |
| $w_{consecutive\ y, thrust}$ | 1/10 |
| $w_{wide\ z, thrust}$ | 3/10 |
| $w_{velocity, lift}$ | 5 |
| $w_{towing, x}$ | 0 |
| $w_{towing, y}$ | 4 |
| $w_{criteria, towing, y}$ | 0 |
| $w_{towing, z}$ | 0 |
| $\vec{w}_{emergence}$ | $\vec{0}$ |

**Table A3:** Weights for the lift force step.

| Weight | Value |
| --- | --- |
| $w_{mass\ factor}$ | 1/50 |
| $g$ | 10 |
| $w_{thrust, mobility}$ | 1/3 |
| $w_{lift, mobility}$ | 1/10 |
| $w_{thermal, total}$ | 1/3 |
| $w_{mobility, total}$ | 2/3 |
| $y_{criteria, thermal}$ | 1 |
| $w_{thermal, thermal}$ | 3 |
| $w_{consecutive\ y, thermal}$ | 5 |
| $w_{else, thermal}$ | 5 |
| $w_{thrust, thrust}$ | 1 |
| $w_{consecutive\ y, thrust}$ | 3/10 |
| $w_{wide\ z, thrust}$ | 6/10 |
| $w_{velocity, lift}$ | 30 |
| $w_{towing, x}$ | 8 |
| $w_{criteria, towing, x}$ | 0 |
| $w_{towing, y}$ | 0 |
| $w_{towing, z}$ | 0 |
| $\vec{w}_{emergence}$ | $\vec{0}$ |

**Figure A1:**
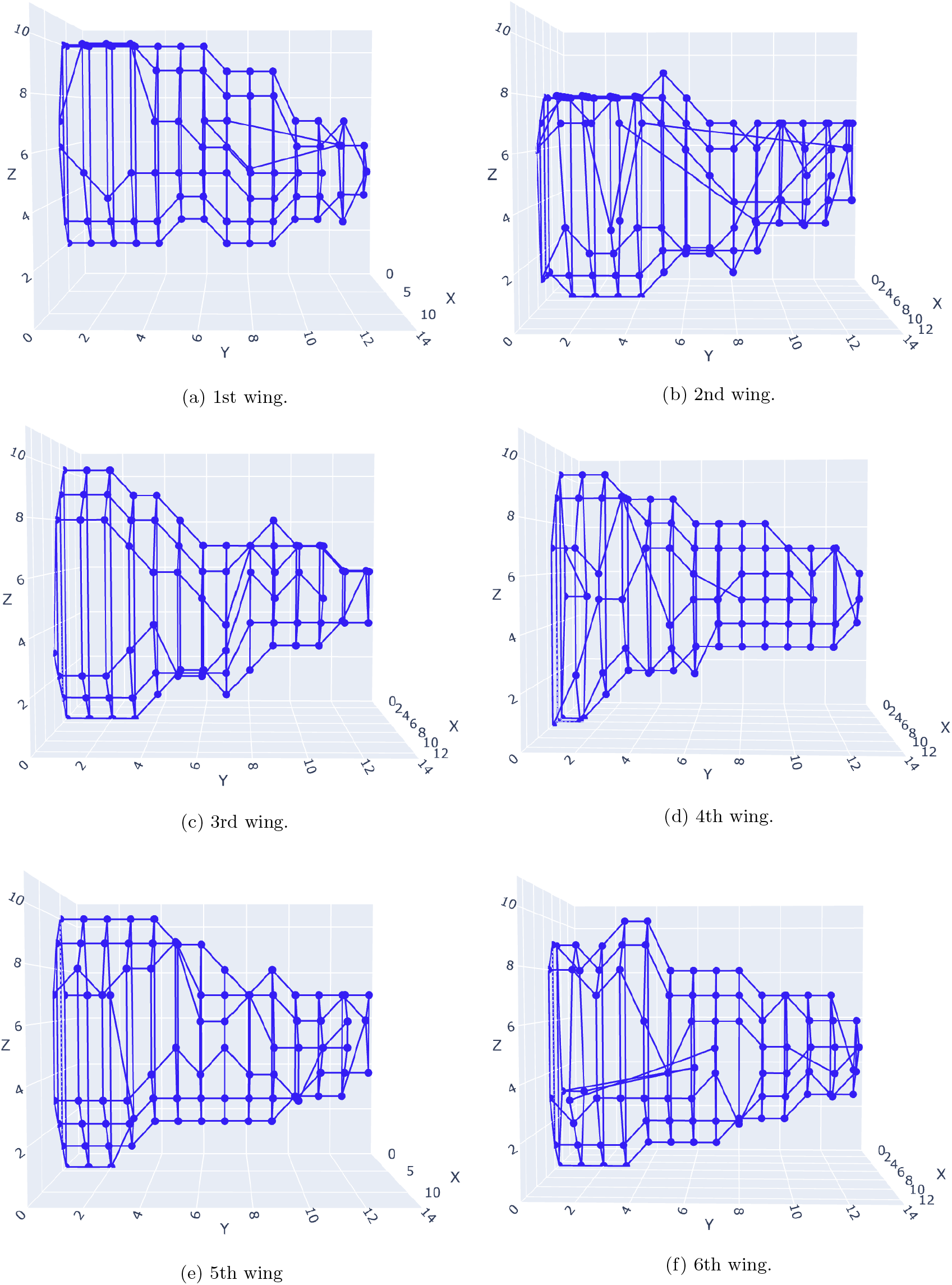
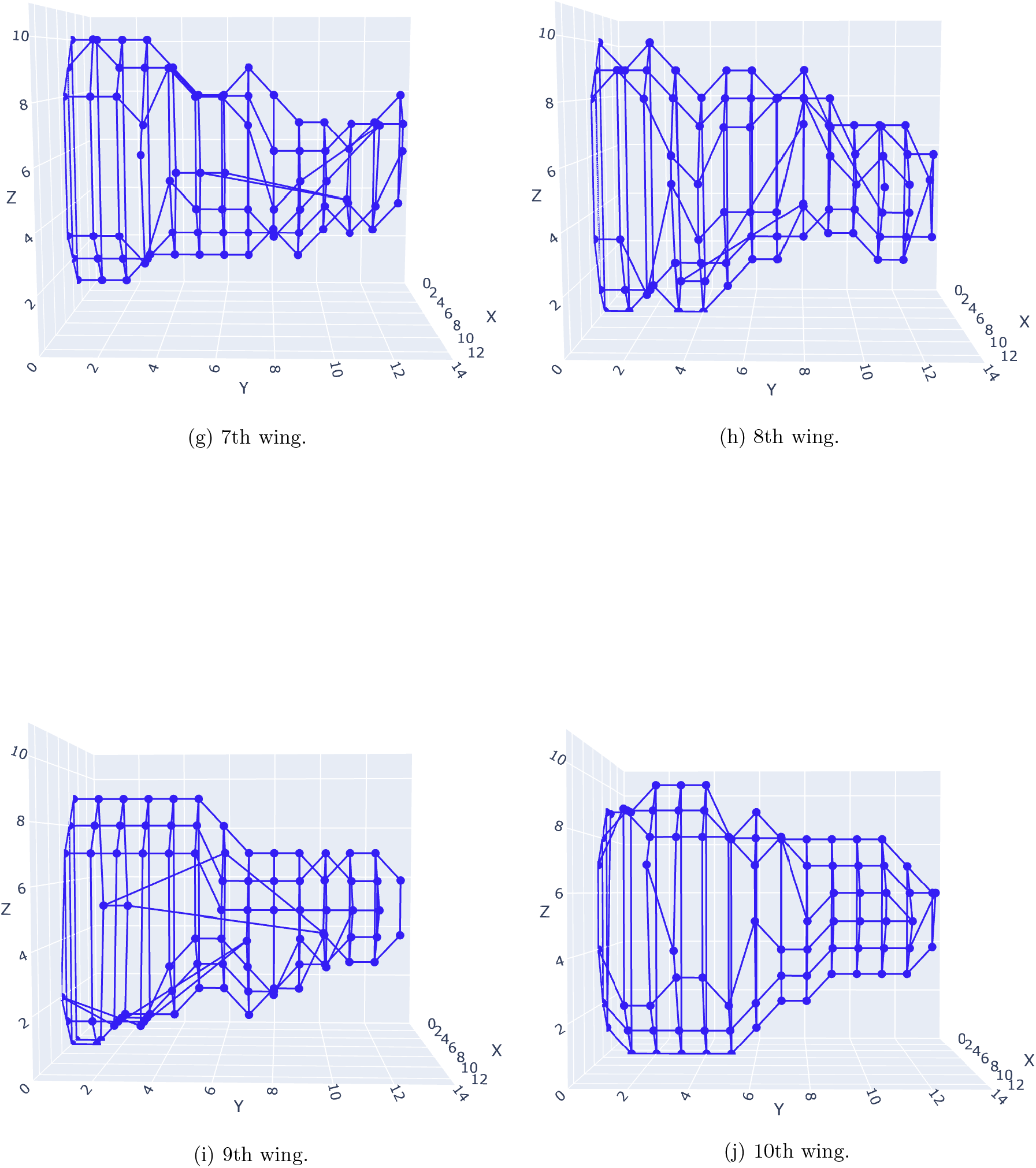
The resulting wing morphologies from DE.

